# Single-cell phylodynamics reveal rapid late-stage colorectal cancer expansions

**DOI:** 10.1101/2025.11.25.690378

**Authors:** Joao M Alves, Kylie Chen, Sonia Prado-López, Nuria Estévez-Gómez, Pilar Alvariño, Juana Alonso, Monica Valecha, Laura Tomás, Antoine Zwaans, Ugnė Stolz, Tanja Stadler, Debora Chantada, José Manuel Cameselle-Teijeiro, Alexei Drummond, David Posada

## Abstract

Single-cell whole-genome sequencing of 335 cells from seven colorectal cancers, coupled with Bayesian phylodynamic modeling, revealed tumors often originate decades before diagnosis, remain indolent, and expand rapidly within two years. Strong spatial structuring with minimal interregional mixing was observed, while substitution rates varied widely and were decoupled from growth. Our findings highlight extended evolutionary stasis and sudden expansion, informing strategies for early detection and intervention.

## Main

Colorectal cancer (CRC) is a leading cause of cancer mortality worldwide^1^. Like many solid tumors, it develops through successive clonal expansions and malignant dissemination^2,3^. While the genetic hallmarks of CRC are well defined^4^, the temporal and spatial dynamics underlying initiation, diversification, and metastasis remain incompletely understood. Genomic studies and phylogenetic modeling have revealed highly variable progression rates, from tumors emerging within years to lineages persisting for decades^3,5–8^, alongside complex, often early, dissemination patterns and diverse routes of clonal migration, underscoring fundamental differences in growth dynamics and ecological interactions ^3,5,8–10^.

At present, most cancer evolutionary studies still rely on bulk sequencing, which averages genomic signals across many cells and requires complex deconvolution, often relying on oversimplified mutational models^11,12^. Single-cell DNA sequencing overcomes these issues by directly profiling individual genomes^13–16^. Still, its adoption is hindered by high costs and technical artifacts, especially allelic dropout (ADO) and whole-genome amplification errors, that can mask true variants and bias phylogenetic reconstruction^17^. Furthermore, most single-cell whole-genome sequencing (scWGS) datasets focus on large-scale copy number variants (CNVs)^18^, whose overlapping nature violates standard phylogenetic assumptions^19^, leaving fine-scale tumor evolution poorly characterized.

Here, we present a high-resolution phylodynamic analysis of 335 single cells from seven CRC patients, sequenced to depths that enable robust genome-wide single-nucleotide variant (SNV) detection. Sampling spanned 31 regions—two to seven anatomically distinct tumor sites per patient—plus matched healthy tissue (**Fig. 1a, Table S1**). Somatic mutation burdens ranged from 4,035 in CRC09 to 11,725 in CRC07, with clonal SNV proportions varying from 18.2% in CRC12 to 37% in CRC14, underscoring substantial inter-patient differences in mutational load and clonal architecture (**Fig. 1b**). CNV analyses revealed pronounced heterogeneity both between and within tumors: CRC01 exhibited broad amplifications on chromosomes 8, 13, and 17, with several metastatic “L-LN” cells showing whole-genome duplication, while CRC08 displayed region-specific profiles from near diploid to chromosome-wide gains on 13 and 20 (**Fig. 1c**).

**Figure 1.**
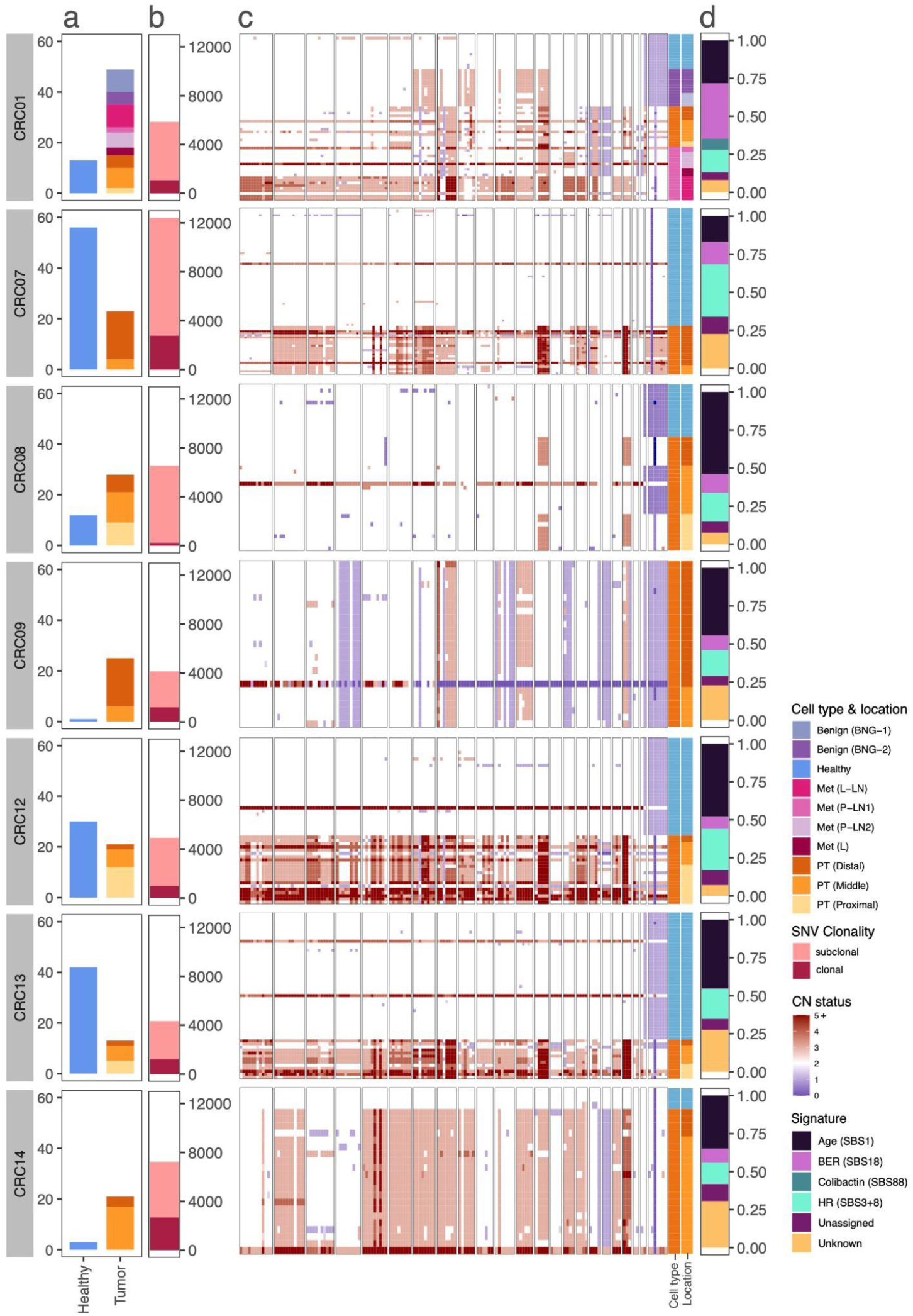
Genomic characterization of single-cell datasets from seven CRC patients. **a.** Stacked barplot depicting the number of single-cells sequenced per patient. Blue = cells from healthy tissue; purple hue = cells from benign tumor; orange hue = cells from primary tumor (PT); pink hue = cells from metastatic lesions (Met): L-LN = hepatic lymph-nodes; P-LN: peripancreatic lymph-nodes; L: Liver. **b**. Bar plots show the number of somatic mutations identified in tumor cells across patients. The different colors highlight clonal (dark pink) and subclonal (light pink) mutations. **c**. Heatmap depicting genome-wide copy-number status (deletions: blue, amplifications: red) of sampled cells. Chromosomes are ordered from 1 to 22 and X. For each panel, the rows represent individual cells. The right-side panels describe the cell type and tissue of origin (using the same color scheme as in **a**.). **d**. Barplots describing the contribution of different mutational signatures: BER = base excision repair; HR = homologous recombination deficiency. SBS signatures with unknown etiology - SBS17, SBS93, SBS94, and SBS121 - were collapsed into a single “Unknown” category.

*De novo* SBS signature extraction identified nine processes, including age-related SBS1, base-excision repair deficiency (SBS18), homologous recombination deficiency (SBS3, SBS8), and the colibactin-linked SBS88, found only in CRC01 (**Fig. 1d**).

Leveraging a novel version of *Phylonco*^*20*^, a state-of-the-art single-cell phylogenetic model within the BEAST2 framework^21^, we reconstructed time-calibrated phylogenies for all CRC patients. Model estimates confirmed the high quality of our scWGS data, with low ADO and error rates (**Supplementary Note 1, Fig. S1**). The inferred trees (**Fig. 2a**) revealed well-supported clades, with cells clustering predominantly by anatomical region, underscoring pronounced spatial structuring in most tumors. Across patients, the number of clonal non-silent mutations ranged from 21 in CRC01 to 61 in CRC14 (**Table S2**), with the majority of alterations in key CRC driver genes (e.g., *APC, TP53, PIK3CA*) being clonal, consistent with early acquisition of critical oncogenic events.

**Figure 2.**
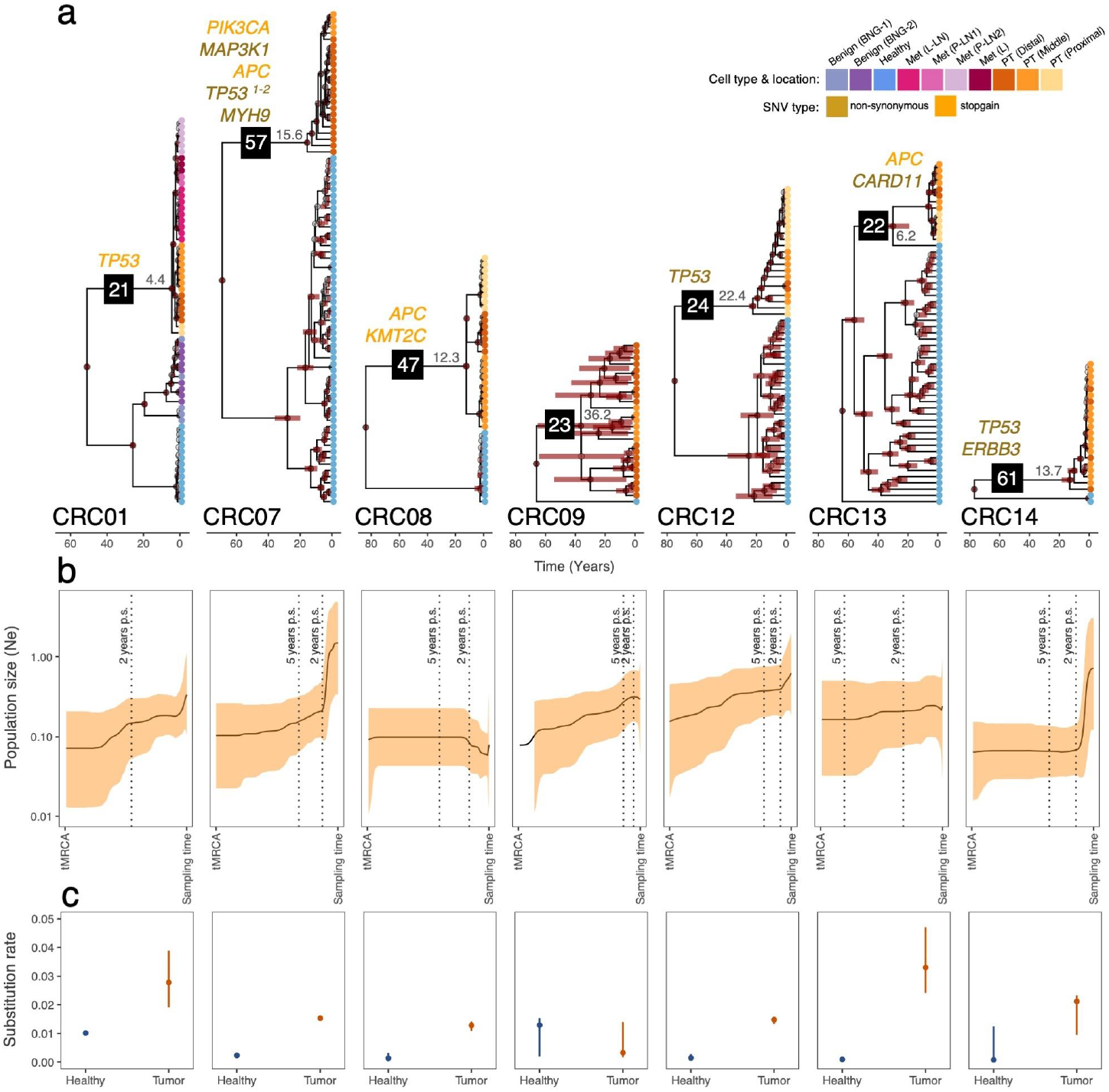
Evolutionary and demographic reconstruction of CRCs over time. **a.** Conditional clade probability distribution (CCD) trees resulting from the *Phylonco* analyses for all patients. Tip colors represent different cell types and sampling locations. Posterior support values above 0.5 are colored according to a continuous transparency scale, with solid black circles representing a posterior value of 1. Red horizontal bars correspond to the 95% highest posterior density (HPD) intervals of the age estimates. Mean estimates of the tumor sample MRCA (tMRCA) are shown in grey. For each patient, the number of non-silent mutations is displayed on the branch leading to the tMRCA (black box), with mutations affecting known CRC driver genes highlighted above. **b**. Bayesian Skyline plot analysis depicting the demographic changes in the tumor cell population over time. For each patient, the solid line represents the historical effective population size (*N*_*e*_) of the tumor, with the shading illustrating the 95% HPD interval. Vertical dashed lines highlight the time at 5 and 2 years before sampling (p.s.). Only one dashed line is shown for patient CRC01 because the tMRCA age estimate is less than 5 years. **c**. Posterior distributions of substitution rates in healthy and tumor cell populations. Solid points depict the posterior median with vertical bars showing the 95% HPD interval.

Importantly, our phylogenetic analyses indicate that several tumors began evolving decades before diagnosis, with a mean time to the most recent common ancestor of the sample (tMRCA) of 15.8 years before diagnosis. This estimate captures only the divergence of the sampled lineages and thus represents a conservative lower bound on tumor age. All patients, except CRC09, exhibited narrow 95% highest posterior density (HPD) intervals, indicating well-constrained tMRCA estimates. In contrast, CRC09 displayed substantial uncertainty, with a broad 95% HPD range spanning 7.4 to 65 years. Among patients with precise tMRCA estimates, CRC12 showed the oldest tMRCA (22.4 years), whereas CRC01 had the most recent estimate (4.4 years) and experienced rapid expansion. In CRC01, all metastatic cells formed a single, well-supported clade, consistent with a monoclonal origin^7^. The metastatic ancestor diverged approximately 1.9 years after the tMRCA, while a benign lesion in the same patient emerged independently about 19 years before sampling, substantially predating the malignant tumor (**Fig. S2**).

We quantified temporal changes in the effective population size (*N*_*e*_) of tumor cell populations using Bayesian skyline plots (Fig. 2b), which revealed distinct demographic trajectories across patients. CRC01, CRC09, and CRC12 showed gradual, sustained growth, consistent with a continuous expansion, while CRC08 and CRC13 remained largely stable after the tMRCA. In CRC07 and CRC14, *N*_*e*_ surged sharply approximately 2 years preceding sampling, following prolonged periods of minimal growth. Although no new mutations were found in established CRC driver genes to account for these late expansions, CRC07 harbored non-synonymous changes in *SERPINE1* and *ADAMTS4*, suggesting possible roles in late-stage growth (**Supplementary Note 2, Fig.S3**)^22,23^.

Substitution rates were consistently higher in tumor cells compared with their healthy counterparts (**Fig. 2c**). Median rates varied markedly across patients, ranging from modest differences (∼3-fold in CRC01) to pronounced disparities (∼35-fold in CRC13). In CRC09 and CRC14, however, the limited number of healthy cells yielded broad 95% HPD intervals that substantially overlapped with those of the tumor estimates, preventing confident resolution of rate differences.

To investigate the spatial dynamics of tumor dissemination, we performed discrete phylogeographic analyses to trace migration events among anatomical regions within each patient. These reconstructions revealed well-defined geographical origins and patient-specific colonization patterns (**Fig. 3**), except in CRC08, where similar posterior probabilities across regions precluded a definitive assignment.

**Figure 3.**
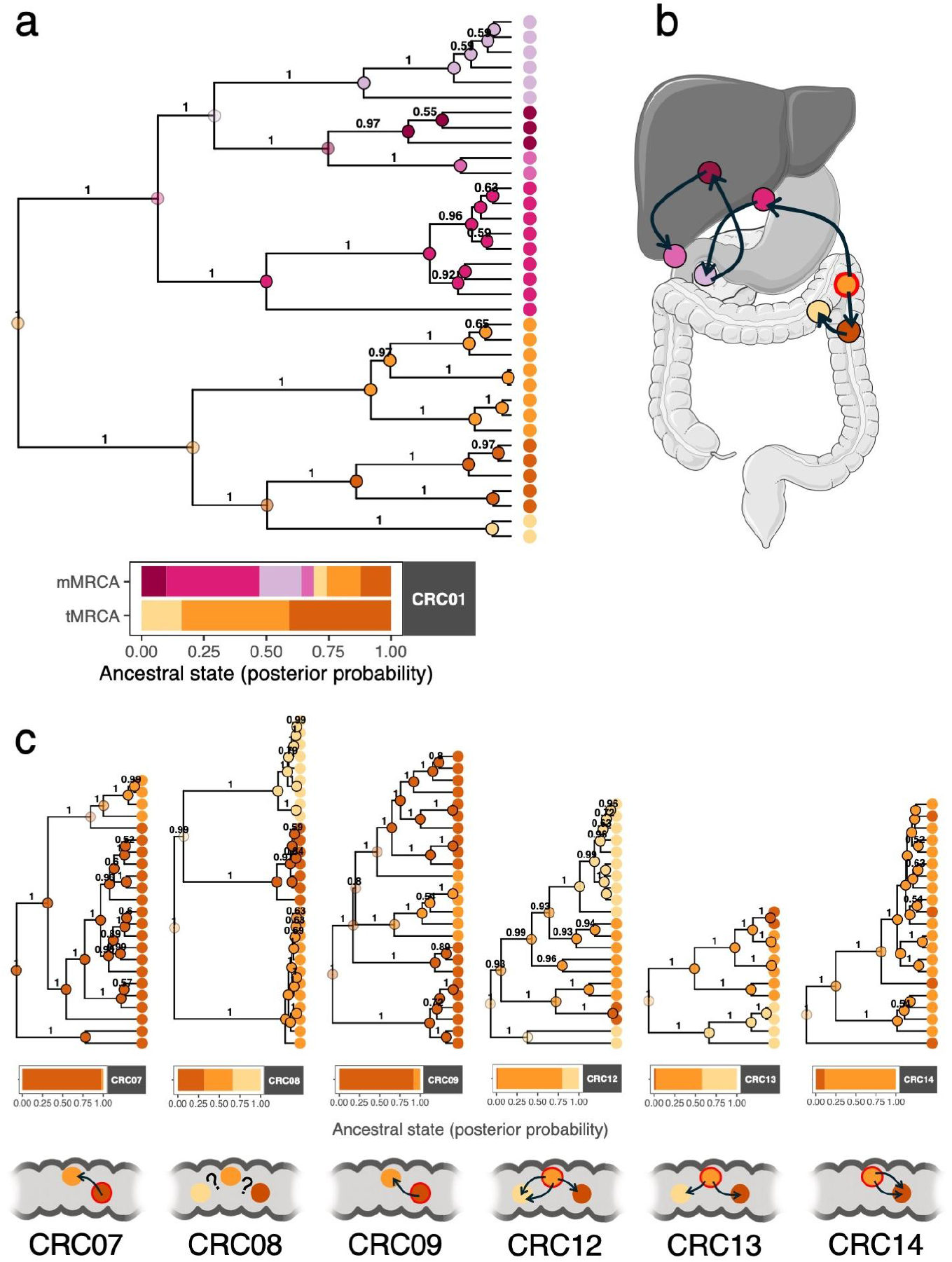
Inferred phylogeographic history from single-cell phylogenies. **a.** Phylogeographic reconstruction of colonization patterns and range expansions of tumor lineages describing the inferred location of all ancestral lineages in patient CRC01. At each tree node, the geographical location with the highest posterior probability was colored using a continuous transparency scale, with solid black circles representing a posterior value of 1.0. Tips and internal nodes are colored according to sampling location. The bar plot below shows the posterior probabilities for each sampled location of the tumor (tMRCA) and metastatic (mMRCA) ancestral lineages. **b**. Schematic representation illustrating malignant cell population dynamics over time in CRC01. Colored circles represent sampled locations with arrows indicating movement direction. Image adapted from Servier Medical Art (https://smart.servier.com/), licensed under CC BY 4.0 (https://creativecommons.org/licenses/by/4.0/). The red outline indicates the inferred tumor origin. **c**. Phylogeographic reconstruction, posterior probability of tMRCA location, and tumor cell population dynamics for the remaining patients. The red outline indicates the inferred tumor origin.

In CRC01, the tMRCA was inferred to originate in the middle section of the primary tumor, with descendant lineages radiating in multiple directions (**Fig. 3a-b**). Although the posterior support was limited, the ancestral metastatic lineage likely underwent an early long-distance migration to the hepatic lymph nodes, followed by sequential spread to the liver and peripancreatic lymph nodes –a trajectory consistent with a stepwise metastatic cascade^24^. Using *Mascot*^*25*^, a structured coalescent model that jointly infers migration and regional demography, we found that metastatic expansion within the liver and peripancreatic nodes accounted for much of the overall tumor growth (**Fig. S4**). This expansion coincided with the mMRCA, suggesting that successful colonization of these niches drove the subsequent proliferation. Mutation mapping along metastatic branches revealed a non-synonymous substitution in *TF*, a gene implicated in promoting hepatic metastasis and CRC invasion^26^ (**Supplementary Note 2, Fig.S3**).

In the remaining patients (**Fig. 3c**), migration patterns supported predominantly localized expansions. In CRC07 and CRC09, tumors originated in the distal region before spreading toward the middle section. CRC12, CRC13, and CRC14 likely originated in the middle section; CRC12 showed multiple migrations toward proximal and distal sites, CRC13 followed a simpler bidirectional route, and CRC14 displayed minimal spatial structure, consistent with continuous cell mixing.

In summary, our study provides one of the most comprehensive scWGS reconstructions of CRC evolution to date. By combining Bayesian phylodynamic and phylogeographic modeling with multiregional sampling, we inferred divergence times, population dynamics, substitution rates, and migration pathways while explicitly accounting for scWGS noise and phylogenetic uncertainty. We found that CRC tumors are strongly spatially structured, with cells clustering by anatomical origin, indicating limited interregional mixing. Migration patterns were generally simple, reflecting stepwise local expansions. Crucially, phylogenetic dating showed that most tumors originated decades before diagnosis^3,6^, endured prolonged indolent phases, and then underwent rapid expansions within roughly two years. However, both the onset and pace of growth varied substantially across patients. Substitution rates were consistently elevated in tumor cells compared to matched healthy cells, yet their magnitude was decoupled from growth tempo. These results highlight extended, low-growth periods that may provide actionable windows for early diagnosis and intervention.

Overall, our work demonstrates the power of single-cell phylodynamics to resolve the tempo and mode of cancer evolution, linking fine-scale genomic patterns to spatial and temporal dynamics. Future integration of these temporal reconstructions with functional and clinical data from larger patient cohorts will be essential for identifying early evolutionary hallmarks of malignancy and translating them into actionable biomarkers.

## Material & Methods

### Sample acquisition and processing

Tissue samples were obtained from seven patients with histologically confirmed colorectal cancer (CRC). For each patient, the samples included the primary tumor, matched metastases (if available), and adjacent normal colon tissue. Samples were collected either during a warm autopsy (patient CRC01) at the University Clinical Hospital of Santiago de Compostela, Spain, or during routine clinical procedures (patients CRC07, CRC08, CRC09, CRC12, CRC13, and CRC14) at the University Hospital Álvaro Cunqueiro, Vigo, Spain. For each patient, 2-3 samples were taken from the primary tumor. In the case of CRC01, additional samples included one liver metastasis, three regional lymph node metastases, and two benign tumor samples. All samples were snap-frozen in liquid nitrogen directly after surgical excision and stored at −80 °C. All tissue samples were obtained from the IDIS-CHUS Biobank (PT13/0010/0068) and the Galicia Sur Health Research Institute (IISGS) Biobank (B.0000802), members of the Spanish National Biobank Network. Informed consent was obtained from all participants before tissue collection. This study was approved by a local Ethical and Scientific Committee (CAEI Galicia 2014/015).

### Tissue disaggregation and single-cell sorting

We minced each sample into 1 mm^3^ pieces with a scalpel and incubated them in Accutase (LINUS) for 1 hour at 37 °C. We filtered the cell suspensions with a 70 μm cell strainer (Falcon, NY, USA) and assessed cell viability with Trypan Blue (Gibco, MA, USA). When the percentage of dead cells exceeded 30%, we performed a Ficoll-Paque density gradient centrifugation to discard dead cells and debris before sorting. We washed the cell pellets twice and resuspended them in ice-cold phosphate-buffered saline (PBS). Then, we incubated them with antibodies against the epithelial cell adhesion molecule (EpCAM) (Clone EBA1, FITC, 347197) and stem cell markers CD44 (APC, 559942), CD166 (PE, 559263), and Lgr5 (BV 421562925) (all from BD Biosciences, NJ, USA). The DRAQ5 (Thermo Fisher Scientific, MA, USA) and 7AAD (Invitrogen, MA, USA) dyes were additionally used to select nucleated cells and exclude non-viable ones. Cells were gated and sorted based on the expression of the chosen markers using a FACSAria III flow cytometer (BD Biosciences, NJ, USA). We analyzed the resulting data using FACSDiva (BD Biosciences, NJ, USA) and FlowLogic software (Miltenyi Biotec, Germany).

### Single-cell whole-genome amplification

To obtain sufficient DNA for sequencing, we performed single-cell whole-genome amplification (scWGA) using the Ampli1 Kit (Menarini Silicon Biosystems, Italy). To avoid contamination, we worked under a biological safety cabinet, used a dedicated set of pipettes, and UV-irradiated all the plastic materials employed. In addition to the patients’ cells, we included a positive control (10 ng/μl REPLIg human control kit, QIAGEN, Netherlands) and a negative control (DNase/RNase-free water) in the amplification process. We assessed the quality of the amplified DNA with the Ampli1 QC Kit (Menarini Silicon Biosystems, Italy). For positive samples for the four Ampli1 QC PCR fragments, we used the Ampli1 ReAmp/ds kit (Menarini Silicon Biosystems, Italy) to increase the total amount of double-stranded DNA. We then removed the Ampli1 adapters, adding 5 μl of NE Buffer 4 10X (New England Biolabs, MA, USA), 1 μl of MseI 50U/μl (New England Biolabs, MA, USA), and 19 μl of nuclease-free water to every 25 μl of sample, and incubating this mixture at 37 °C for 3 h, followed by 20 min at 65 °C for enzyme inactivation. After incubation, we purified the samples using 1.8X AMPure XP beads (Agencourt, Beckman Coulter, CA, USA). Finally, we quantified the DNA yield using a Qubit 3.0 fluorometer (Thermo Fisher Scientific, MA, USA) and checked the amplicon size distribution with the D5000 ScreenTape assay in a 2200 TapeStation platform (Agilent Technologies, CA, USA).

### Bulk genomic DNA (gDNA) isolation

For each patient, we additionally isolated the gDNA from a normal bulk sample using the QIAamp DNA Mini kit (QIAGEN, Netherlands). Next, we estimated DNA yield using the Qubit 3.0 fluorometer (Thermo Fisher Scientific, MA, USA) and DNA integrity using the Genomic DNA ScreenTape assay on the 2200 TapeStation platform (Agilent Technologies, CA, USA).

### Library preparation and sequencing

Three hundred and thirty-five single-cell and seven bulk samples were submitted to the Spanish National Center for Genomic Analysis (CNAG; http://www.cnag.crg.eu) for library construction and sequencing. Sequencing libraries were constructed with the KAPA HyperPrep kit (Roche, Sweden). Sequencing was performed on an Illumina NovaSeq 6000 platform (PE150), with an expected depth of approximately 6X for the single-cell libraries and 30X for the bulk samples.

### Pre-processing and variant calling

We aligned the sequencing reads from both single-cell and bulk DNA libraries to the Genome Reference Consortium Human Build 37 (GRCh37) using the BWA-MEM algorithm^27^. A standardized best-practices pipeline^28^ was followed, including filtering of low-quality reads, local realignment around indels, and removal of PCR duplicates. We identified somatic single-nucleotide variants (SNVs) in individual cells using SCcaller^29^, a variant caller specifically tailored for single-cell DNA sequencing data, under default parameters. Following variant detection, VCFs for each patient were merged using bcftools^30^. To ensure high-confidence variant calls, we applied stringent filtering criteria, retaining only SNVs annotated as “True” somatic mutations and observed in at least two individual single cells. Finally, SNV annotations were performed using ANNOVAR (version 20200608)^31^. In parallel, we inferred single-cell copy number variants (CNVs) using GINKGO^32^ with approximately 500-kb variable-length bins. CNV data were normalized and segmented per cell using default settings.

### Mutational signatures

We ran the SIGNAL web tool^33^, 2020) with default parameters to identify single-base substitution (SBS) signatures active in tumor cells from each patient. Following the tool’s recommendation, SBS fitting was performed using candidate SBS signatures from CRC.

### Phylogenetic reconstruction and dating

We used BEAST 2.7^21^ with the Phylonco v1.2.1^20^ and FlexibleLocalClock (FLC) v1.2.0 packages^34^, and Beagle^35^ with GPU acceleration to run the analyses. We used a GT16 substitution model with a Dirichlet(3,3,…,3) prior on frequencies and a Dirichlet(1,2,1,1,2,1) prior on the relative rates. Error parameters were estimated using a GT16 error model^20,36^ with a Beta(1, 2) prior on allelic dropout and the combined sequencing/amplification error. Local clocks were used to account for differences in clock rates between healthy and tumor cells. We defined one strict clock for healthy cells and another for tumor cells, both with a Lognormal prior (mean=-5.5, sd=1.65) on the clock rate. To account for population growth, we used a Bayesian skyline coalescent model^37^ with a Lognormal prior (mean=2.35, sd=2.35) on population size. The tumor cells were constrained to be monophyletic in all analyses. For each dataset, we applied a time calibration spanning a 1-year window around the patient’s age at the root of the phylogeny. This assumes that the divergence time between the healthy and tumor lineages occurred around birth. Our sampling design and biological evidence support this assumption. Our healthy and tumor cells were obtained from spatially distinct regions, corresponding to independent colonic crypts, and prior work has shown that crypts are long-lived, clonally independent units established during early development^38^. Two independent Monte Carlo Markov Chains (MCMC) were run for each patient and combined after removing the first 10% of the chain (burn-in). The combined chains had between 150 million and 1,000 million samples. The effective sample size (ESS) of the identifiable parameters were over 200, except for CRC13, which has a bimodal estimate of the tMRCA, reducing the ESS (in CRC13, a burn-in of 70% was used). The posterior trees were summarized using the conditional clade distribution metric CCD0^39^.

### Demographic analysis

Tumor demographics were reconstructed using a Bayesian skyline coalescent approach^37^ on the tumor cell population using a Lognormal prior (mean=2.35, sd=2.35) on the population size. The estimated effective population size was scaled using the mean tMRCA time from the reconstructed phylogenies.

### Phylogeography and migration history

We conducted a discrete phylogeography analysis of the tumor cell population using the beast-classic v1.6.3 package^40^ in BEAST 2.7, with the demographic model described above. The primary tumor and metastatic regions were used as locations for the geographic model, with a Gamma(1,1) prior on relative migration rates, and assuming that the tMRCA originated in one of the primary tumour samples. In addition, a structured coalescent analysis was run using BEAST’s package *Mascot* v3.0.7, allowing for differential population growth among regions from patient CRC01^25^, with an Exponential(1) prior on migration rates between areas. In each region, a Normal(0,1) prior was used for the relative changes through time in effective population size, whose dynamics follow a Gaussian Markov Random Field smoothing assumption^41^. The estimated effective population sizes were scaled using the mean tMRCA time from the reconstructed phylogenies. Both the phylogeography and migration analyses used the same substitution model as the unstructured Bayesian skyline analysis described above. For both the discrete phylogeography and Mascot analyses, two independent MCMC chains of at most 100 million samples were run and combined after removing a burn-in of 10%. The ESS of identifiable parameters in the combined results was over 200 in all cases. The posterior trees were summarized using the conditional clade distribution metric CCD0.

## Code availability

Phylonco software is available at https://github.com/bioDS/beast-phylonco. All code and analyses for this study are available on GitHub at https://github.com/kche309/phylocancer.

## Data availability

We deposited the raw single-cell and bulk whole-genome sequencing data for all patients at the National Center for Biotechnology Information (NCBI) as BioProject ######.

## Author contributions

DP conceived and supervised the study. JMC-T and DC obtained the tumor samples. SPL, NEG, PA, and JA processed the samples. JMA, KC, MV, AZ, LT, and US performed the computational analyses. AJD and TS contributed to the conceptualisation of the models and technical decisions related to the phylodynamic and phylogeographic analyses. JMA and DP wrote the manuscript with input from all other authors.

## Acknowledgments

We want to thank the Supercomputation Center of Galicia (CESGA) for providing computational resources.

## References

1. Bray, F. et al. Global cancer statistics 2022: GLOBOCAN estimates of incidence and mortality worldwide for 36 cancers in 185 countries. CA Cancer J. Clin. 74, 229–263 (2024).

2. Merlo, L. M. F., Pepper, J. W., Reid, B. J. & Maley, C. C. Cancer as an evolutionary and ecological process. Nature Reviews Cancer 6, 924–935 (2006).

3. Gerstung, M. et al. The evolutionary history of 2,658 cancers. Nature 578, 122–128 (2020).

4. Li, J., Ma, X., Chakravarti, D., Shalapour, S. & DePinho, R. A. Genetic and biological hallmarks of colorectal cancer. Genes Dev. 35, 787–820 (2021).

5. Zhao, Z.-M. et al. Early and multiple origins of metastatic lineages within primary tumors. Proc. Natl. Acad. Sci. U. S. A. 113, 2140–2145 (2016).

6. Lote, H. et al. Carbon dating cancer: defining the chronology of metastatic progression in colorectal cancer. Ann. Oncol. 28, 1243–1249 (2017).

7. Alves, J. M., Prado-López, S., Cameselle-Teijeiro, J. M. & Posada, D. Rapid evolution and biogeographic spread in a colorectal cancer. Nat Commun 10, 5139 (2019).

8. Gabbutt, C. et al. Fluctuating DNA methylation tracks cancer evolution at clinical scale. Nature (2025) doi:10.1038/s41586-025-09374-4.

9. Hu, Z. et al. Quantitative evidence for early metastatic seeding in colorectal cancer. Nat. Genet. 51, 1113–1122 (2019).

10. Welch, H. G. & Black, W. C. Overdiagnosis in cancer. J. Natl. Cancer Inst. 102, 605–613 (2010).

11. Tanner, G., Westhead, D. R., Droop, A. & Stead, L. F. Benchmarking pipelines for subclonal deconvolution of bulk tumour sequencing data. Nat. Commun. 12, 6396 (2021).

12. Salcedo, A. et al. Crowd-sourced benchmarking of single-sample tumor subclonal reconstruction. Nat. Biotechnol. 43, 581–592 (2025).

13. Baslan, T. & Hicks, J. Unravelling biology and shifting paradigms in cancer with single-cell sequencing. Nat Rev Cancer 17, 557–569 (2017).

14. Roth, A. et al. Clonal genotype and population structure inference from single-cell tumor sequencing. Nat. Methods 13, 573–576 (2016).

15. Leung, M. L. et al. Single-cell DNA sequencing reveals a late-dissemination model in metastatic colorectal cancer. Genome Res. 27, 1287–1299 (2017).

16. Alves, J. M. et al. Clonality and timing of relapsing colorectal cancer metastasis revealed through whole-genome single-cell sequencing. Cancer Lett 543, 215767 (2022).

17. Estévez-Gómez, N. et al. Differential performance of strategies for single-cell whole-genome amplification. Cell Rep. Methods 5, 101025 (2025).

18. McPherson, A. et al. Ongoing genome doubling shapes evolvability and immunity in ovarian cancer. Nature 644, 1078–1087 (2025).

19. Zaccaria, S., El-Kebir, M., Klau, G. W. & Raphael, B. J. Phylogenetic copy-number factorization of multiple tumor samples. J. Comput. Biol. 25, 689–708 (2018).

20. Chen, K., Moravec, J. C., Gavryushkin, A., Welch, D. & Drummond, A. J. Accounting for Errors in Data Improves Divergence Time Estimates in Single-cell Cancer Evolution. Mol Biol Evol 39, (2022).

21. Bouckaert, R. et al. BEAST 2.5: An advanced software platform for Bayesian evolutionary analysis. PLoS Comput. Biol. 15, e1006650 (2019).

22. Wang, Y., Wang, J., Gao, J., Ding, M. & Li, H. The expression of SERPINE1 in colon cancer and its regulatory network and prognostic value. BMC Gastroenterol. 23, 33 (2023).

23. Chen, J. et al. Promotion of tumor growth by ADAMTS4 in colorectal cancer: Focused on macrophages. Cell. Physiol. Biochem. 46, 1693–1703 (2018).

24. Turajlic, S. & Swanton, C. Metastasis as an evolutionary process. Science 352, 169–175 (2016).

25. Müller, N. F., Bouckaert, R. R.Wu, C.-H. & Bedford, T. MASCOT-Skyline integrates population and migration dynamics to enhance phylogeographic reconstructions. PLoS Comput. Biol. 21, e1013421 (2025).

26. Tian, M. et al. Depletion of tissue factor suppresses hepatic metastasis and tumor growth in colorectal cancer via the downregulation of MMPs and the induction of autophagy and apoptosis. Cancer Biol. Ther. 12, 896–907 (2011).

27. Li, H. Aligning sequence reads, clone sequences and assembly contigs with BWA-MEM. arXiv [q-bio.GN] (2013) doi:10.48550/ARXIV.1303.3997.

28. Van der Auwera, G. A. et al. From FastQ data to high-confidence variant calls: The Genome Analysis Toolkit best practices pipeline. Curr. Protoc. Bioinformatics 43, (2013).

29. Dong, X. et al. Accurate identification of single-nucleotide variants in whole-genome-amplified single cells. Nat. Methods 14, 491–493 (2017).

30. Danecek, P. et al. Twelve years of SAMtools and BCFtools. Gigascience 10, (2021).

31. Wang, K., Li, M. & Hakonarson, H. ANNOVAR: functional annotation of genetic variants from high-throughput sequencing data. Nucleic Acids Res. 38, e164 (2010).

32. Garvin, T. et al. Interactive analysis and assessment of single-cell copy-number variations. Nat. Methods 12, 1058–1060 (2015).

33. Degasperi, A. et al. A practical framework and online tool for mutational signature analyses show inter-tissue variation and driver dependencies. Nat. Cancer 1, 249–263 (2020).

34. Fourment, M. & Darling, A. E. Local and relaxed clocks: the best of both worlds. PeerJ 6, e5140 (2018).

35. Ayres, D. L. et al. BEAGLE: an application programming interface and high-performance computing library for statistical phylogenetics. Syst. Biol. 61, 170–173 (2012).

36. Kozlov, A., Alves, J. M., Stamatakis, A. & Posada, D. CellPhy: accurate and fast probabilistic inference of single-cell phylogenies from scDNA-seq data. Genome Biol. 23, 37 (2022).

37. Drummond, A. J., Rambaut, A., Shapiro, B. & Pybus, O. G. Bayesian coalescent inference of past population dynamics from molecular sequences. Mol. Biol. Evol. 22, 1185–1192 (2005).

38. Lee-Six, H. et al. The landscape of somatic mutation in normal colorectal epithelial cells. Nature 574, 532–537 (2019).

39. Berling, L. et al. Accurate Bayesian phylogenetic point estimation using a tree distribution parameterized by clade probabilities. PLoS Comput. Biol. 21, e1012789 (2025).

40. Lemey, P., Rambaut, A., Drummond, A. J. & Suchard, M. A. Bayesian phylogeography finds its roots. PLoS Comput. Biol. 5, e1000520 (2009).

41. Gill, M. S. et al. Improving Bayesian population dynamics inference: a coalescent-based model for multiple loci. Mol. Biol. Evol. 30, 713–724 (2013).

