## Supplementary info for "Single-cell phylodynamics reveal rapid late-stage colorectal cancer expansions"

a. CINBIO, Universidade de Vigo, 36310 Vigo, Spain
b. Galicia Sur Health Research Institute (IIS Galicia Sur), SERGAS-UVIGO, Spain
c. School of Biological Sciences, University of Auckland, Auckland, New Zealand
d. Department of Biosystems Science and Engineering, ETH Zürich, Basel, Switzerland
e. Swiss Institute of Bioinformatics (SIB), Lausanne, Switzerland
f. Department of Pathology, Hospital Álvaro Cunqueiro, Vigo, Spain
g.Department of Pathology, Clinical University Hospital, Galician Healthcare Service (SERGAS), Santiago de Compostela, Spain
h. Medical Faculty, University of Santiago de Compostela, Santiago de Compostela, Spain
i. School of Computer Science, University of Auckland, Auckland, New Zealand
^ Current address, Research Unit of Nanoelectronic Devices, Institute of Solid State Electronics, Faculty of Electrical Engineering and Information Technology, TU Wien, Vienna, Austria
[illegible] Current address: Centro de Biomedicina Experimental (CEBEGA), Universidade de Santiago de Compostela, 15706 Santiago de Compostela, Spain
§ Current address: Department of Biosystems Science and Engineering, ETH Zürich, Basel, Switzerland

#### Supplementary note

1. **Inference of critical technical parameters from scWGS data with *Phylonco*:**

Accurate phylogenetic reconstruction from scWGS data critically depends on properly accounting for technical artifacts introduced during whole-genome amplification (Navin 2014). Errors such as allelic dropout (ADO) and amplification/sequencing noise can distort variant calls, bias branch lengths, and ultimately affect evolutionary parameter estimates. To mitigate these effects, we used an observational model in Phylonco that jointly estimates technical and evolutionary parameters (Kozlov et al. 2022), allowing us to quantify artifact impacts while ensuring robust inference. Across patients, inferred ADO rates ranged from 0.22 in CRC12 to 0.31 in CRC01, while per-site error rates varied nearly tenfold, from $1.3 \times 10^{-5}$ in CRC07 to $8.8 \times 10^{-5}$ in CRC09 (**Fig. S1**). Despite this variability, error rates remained uniformly low, confirming the overall high quality and reliability of our single-cell datasets

2. **Phylogenetic mapping of mutations associated with recent clonal expansions:**

Given the evidence of recent expansions in some patients (**Fig. 2b**), we sought to pinpoint mutations that may have coincided with or contributed to these demographic events. We mapped non-silent SNVs onto the time-calibrated phylogenies inferred by *Phylonco* using *CellPhy (Kozlov et al. 2022)* to determine the most likely branch placement based on genotype likelihoods and branch lengths.

This analysis focused on CRC01, CRC07, and CRC14, which displayed the most pronounced or recent expansions in $N_e$ (**Fig. S2**). In CRC07 and CRC14, no additional mutations in established CRC driver genes were detected that could explain the sharp late expansion happening around two years before diagnosis. In CRC07, however, we identified two non-synonymous changes in *SERPINE1* and *ADAMTS4*, genes previously linked to CRC progression and invasion, suggesting possible functional roles in late-stage tumor growth (Wang et al. 2023; Chen et al. 2018).

For CRC01, the *Mascot* results indicated that growth was largely driven by metastatic populations in the liver and peripancreatic lymph nodes. Mutation mapping along these metastatic branches revealed a non-synonymous substitution in *TF*, a gene associated with hepatic and lymph node metastasis in CRC (Tian et al. 2011), potentially facilitating successful colonization of these sites.

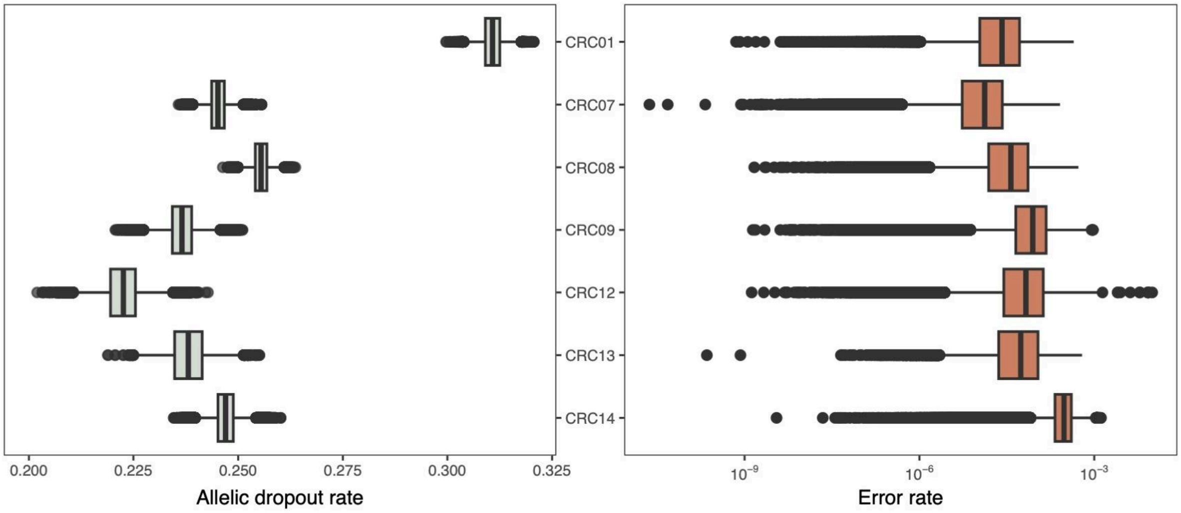


**Figure S1. Estimates of allelic dropout (ADO) and amplification error rates in scWGS datasets.** Box plots show the posterior distributions of the ADO rate (left) and amplification error rate (right) across patients.

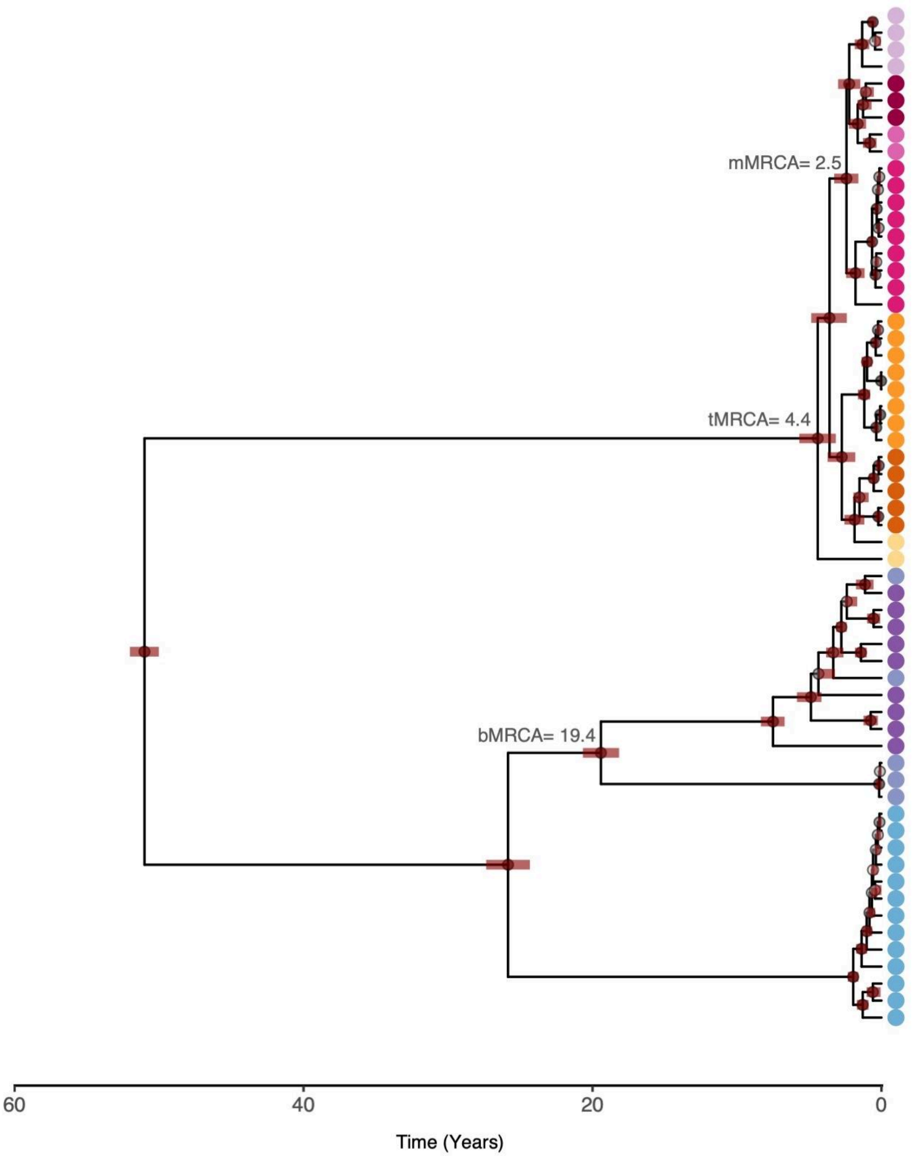


**Figure S2. Single-cell phylogeny of CRC01.** Conditional clade probability distribution (CCD) tree resulting from the *Phylonco* analyses for patient CRC01. Tip colors represent different cell types and sampling locations (using the same color scheme as in Figure 2a). Posterior support values above 0.5 are colored according to a continuous transparency scale, with solid black circles representing a posterior value of 1. Red horizontal bars correspond to the 95% highest posterior density (HPD) intervals of the age estimates. Mean estimates of the age of the primary tumor MRCA (tMRCA), metastasis MRCA (mMRCA), and benign tumor MRCA (bMRCA) are shown in grey.

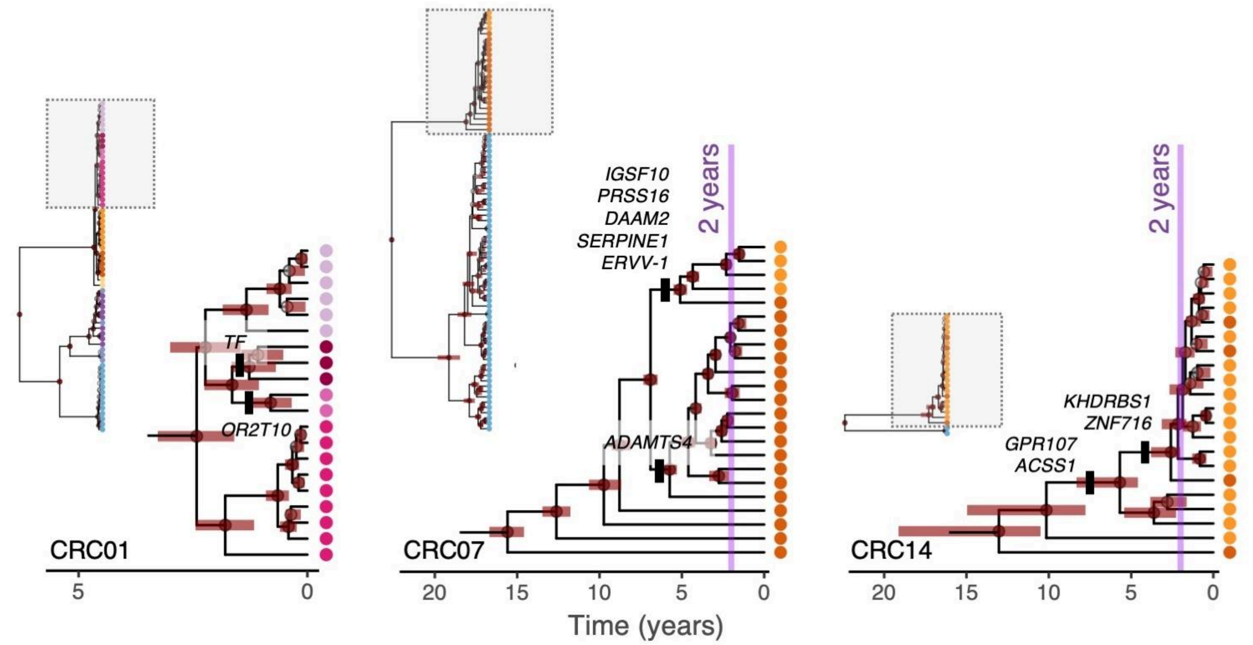


**Figure S3. Temporal mapping of non-silent mutations on the expanding subtrees.** Time-calibrated subtrees for CRC01, CRC07, and CRC14, highlighting the branches corresponding to recent clonal expansions (as inferred from Bayesian skyline plots (Fig. 2b) and the Mascot results (Fig. S4)). For each patient, the whole tree is shown on the left, with the dotted grey square highlighting the portion of the tree evaluated. Gene names indicate the location of mapped non-silent SNVs. Vertical purple lines denote the approximate onset of rapid expansions (approximately two years before sampling) in CRC07 and CRC14. Tip colors represent the different cell types and sampling locations (using the same color scheme as in Figure 2a).

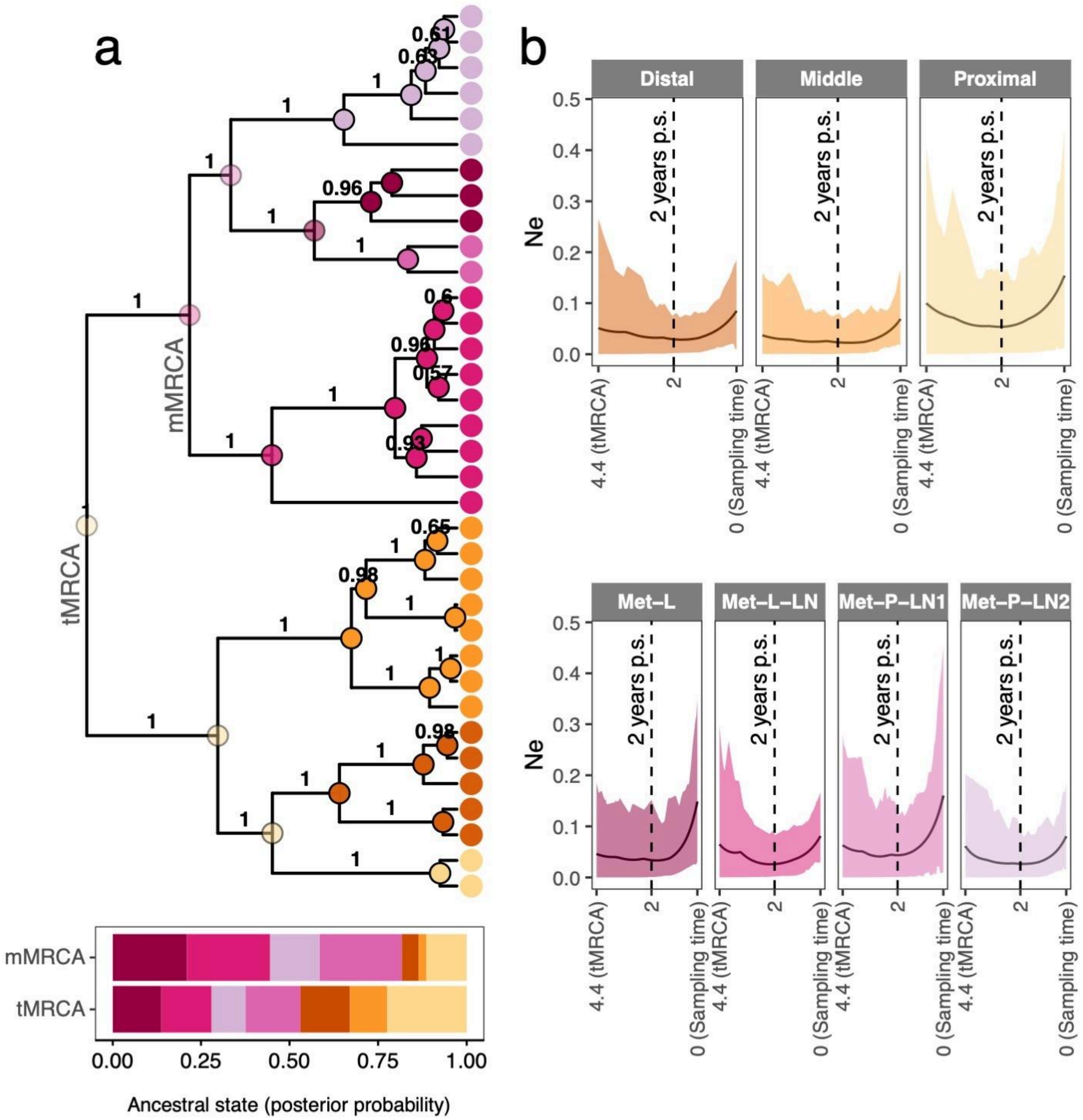


**Figure S4. Phylogeographic history of patient CRC01 using *Mascot*. a.** Bayesian phylogeographic reconstruction of colonization patterns of tumor lineages describing the inferred location of all ancestral lineages. At each tree node, the geographical location with the highest posterior probability was colored using a continuous transparency scale, with solid circles representing a posterior value of 1.0. The barplot at the bottom of the phylogeny shows the posterior probabilities for each sampled location of the tumor (tMRCA) and metastatic (mMRCA) ancestral lineages. **b.** Bayesian Skyline plot analysis depicting demographic changes over time for each sampled region (scaled to the tumour ages estimated in Figure 2). Solid lines represent the historical effective population size ($N_e$) at each site, with the shading illustrating the 95% HPD interval. Vertical dashed lines highlight the time at 2 years prior to sampling (p.s.).

### References


Chen, Jianjun, Yang Luo, Yong Zhou, et al. 2018. "Promotion of Tumor Growth by ADAMTS4 in Colorectal Cancer: Focused on Macrophages." *Cellular Physiology and Biochemistry: International Journal of Experimental Cellular Physiology, Biochemistry, and Pharmacology* 46 (4): 1693–1703.

Kozlov, Alexey, Joao M. Alves, Alexandros Stamatakis, and David Posada. 2022. "CellPhy: Accurate and Fast Probabilistic Inference of Single-Cell Phylogenies from scDNA-Seq Data." *Genome Biology* 23 (1): 37.

Navin, Nicholas E. 2014. "Cancer Genomics: One Cell at a Time." *Genome Biology* 15 (8): 452.

Tian, Maolin, Yuanlian Wan, Jianqiang Tang, et al. 2011. "Depletion of Tissue Factor Suppresses Hepatic Metastasis and Tumor Growth in Colorectal Cancer via the Downregulation of MMPs and the Induction of Autophagy and Apoptosis." *Cancer Biology & Therapy* 12 (10): 896–907.

Wang, Yigang, Jinyan Wang, Jianchao Gao, Mei Ding, and Hua Li. 2023. "The Expression of SERPINE1 in Colon Cancer and Its Regulatory Network and Prognostic Value." *BMC Gastroenterology* 23 (1): 33.
